# Immune regulation of neuronal stem cells in the medaka retina

**DOI:** 10.1101/2025.04.01.646524

**Authors:** Rashi Agarwal, Joergen Benjaminsen, Katharina Lust, Clara Becker, Natalia Fuchs, Eva Hasel de Carvalho, Fanny Eggeler, Omnia El Said Ibrahim, Narges Aghaallaei, Baubak Bajoghli, Joachim Wittbrodt

## Abstract

Stem cell populations in tissues require precise regulation of their number and quality to maintain proper organ growth. Amongst various regulatory mechanisms, immune cells are emerging to directly regulate stem cell populations. The medaka retinal stem cell (RSC) niche, a model for lifelong neurogenic growth, provides a system to study immune-stem cell interactions. We investigate how microglia, resident macrophages of the central nervous system, regulate the RSC niche. We identify that *bona fide* RSCs express the chemokine Ccl25b while its cognate receptor, Ccr9a, is expressed in microglia. These microglia form a surveillance ring adjacent to the RSC niche and actively phagocytose RSCs. Interference with microglia by deletion of *spi1b* reveals that microglia absence leads to increased numbers of ccl25b-positive RSCs and results in morphological defects of the retina. Targeted mutation of *ccl25b* specifically affects microglia mobility under injury conditions, however, we did not observe any morphological defects indicating that Ccl25b-Ccr9a signaling is not essential for stem cell maintenance. Overall, our data show that under homeostatic conditions the individual RSCs, essential for proper eye development, are actively phagocytosed by immune surveillance.

## Introduction

Stem cell reservoirs are essential for growth, homeostasis, and tissue regeneration in multicellular organisms (Hollyfield, 1971; Johns, 1977; Johns & Easter, 1977). These long-lived cells continuously produce new stem and progenitor cells while resisting cell death (Soteriou & Fuchs, 2018). Their quantity and quality must be precisely regulated to prevent tissue overgrowth and cellular deterioration.

The retinal stem cell (RSC) niche of the teleost medaka (Oryzias latipes) provides a well-established model for studying neuronal stem cells (Centanin et al., 2011) due to medaka’s short generation time, ease of genetic manipulation and vast landscape of available transgenic reporter lines. Located in the ciliary marginal zone (CMZ) in a ring around the distal edge of the optic cup, multipotent stem cells sustain lifelong retinal growth (Centanin et al., 2014; Reinhardt et al., 2015; Soteriou & Fuchs, 2018). A population of neuroretinal progenitor cells, located between the stem cell ring and the stratified retina, contains transiently amplifying cells that contribute to both the inner and outer nuclear layers (Tsingos et al., 2019). This distinctive architecture allows reliable identification of multipotent stem cells through lineage tracing experiments (Centanin et al., 2014; Centanin et al., 2011).

Recent evidence suggests that immune cells, particularly tissue-resident macrophages and microglia, extend beyond their classical immune functions to contribute to tissue homeostasis, stem cell regulation, and regeneration (Cunningham et al., 2013; de Almeida et al., 2023; Nath et al., 2024; Ratnayake et al., 2021). In the central nervous system (CNS), microglia are unique among retinal cells as they originate outside the RSC niche, entering the CNS at embryonic stages (E11.5 in mice, 25 hours post fertilization in zebrafish) (Herbomel et al., 2001; Santos et al., 2008).

Traditionally, microglia are known primarily for efferocytosis - the removal of apoptotic cells (Ravichandran, 2011) – and thus contribute to shaping developing tissues. Recently, a “groom or doom” interaction was reported for the hematopoietic stem cell niche of zebrafish and mice during embryogenesis: local macrophages were observed to undergo lengthy interactions with stem cells preceding cell division. These interactions often resulted in either the partial, or complete, phagocytosis of stem cells which did not display markers for apoptosis (Wattrus et al., 2022).

Building on our understanding of the RSC niche organization, we addressed if and how microglia regulate retinal stem cell numbers and contribute to coordinated retinal growth in medaka. Using a newly established retinal stem cell reporter line, we characterize the interactions between RSCs expressing the chemokine ccl25b and microglia expressing its cognate receptor ccr9a.

Unlike the previously established role of ccl25b-ccr9a signaling in the recruitment of lymphocytes in the intestine in mammals and fish (Aghaallaei et al., 2022; Bajoghli, 2013; Uehara et al., 2002), we report a novel function of this chemokine-receptor interaction in the retina. Furthermore, we examine how the complete absence of microglia affects retinal development and growth, providing insights into immune cell-mediated regulation of the retinal stem cell pool.

## Results

### *Ccl25b* chemokine and ccr9a receptor expression in the medaka retina

RSCs are specifically labelled in a stable transgenic line where a ccl25b regulatory element drives GFP reporter expression (*ccl25b::GFP*). This line was initially generated to follow the dynamics of cells expressing *ccl25b*, a chemokine which is known to be involved in the recruitment of lymphocytes in the intestine (Aghaallaei et al., 2022; Campbell & Butcher, 2002; Sigmundsdottir & Butcher, 2008) and thymus (Baubak Bajoghli, 2009). We detected the expression of *ccl25b* chemokine at the periphery of the ciliary marginal zone (CMZ), partially overlapping with previously described *rx2* expressing retinal stem and progenitor cells (Fig. 1A) (Reinhardt et al., 2015). The endogenous expression of *ccl25b* transcripts in the RSC zone was validated by *in situ* hybridization chain reaction (HCR) (Fig.S1B). To test if *ccl25b* expressing cells indeed represent multipotent retinal stem cells, we used GaudíRSG, an established Cre-LoxP based lineage tracing tool (Centanin et al., 2014). Here, Cre induced recombination will lead to the expression of GFP after the Cre mediated excision of a default DsRed/Stop cassette. We employed the ccl25b regulatory element mentioned above (Aghaallaei et al., 2022) to specifically drive the expression of a hormone inducible Cre recombinase in *ccl25b* expressing cells and established a stable transgenic line in the background of the *GaudíRSG* line. We generated a tamoxifen-inducible *ccl25b::Cre^ERT2^,GaudíRSG* transgenic line and clonally activated colour switches in individual *ccl25b*-positive cells specifically labelling those and all of their descendants. The long-term lineage of *ccl25b*-expressing cells was analysed in fish 6-week post Cre-mediated recombination at hatching stage, GFP expressing clones originating from *ccl25b* expressing cells in the CMZ were analysed via wholemount immunostaining. Lineage analysis revealed that *ccl25b*-positive cells established induced <u>Ar</u>ched <u>Co</u>ntinuous <u>S</u>tripes, iArCoSs (Centanin et al., 2014), clonal strings originating in the stem cell niche of the CMZ differentiated into all neuroretinal cell types and thus represent neuroretinal stem cells (Fig. 1B). Early induction for 3 days post hatching was also evaluated to validate the clonal capacity of these ccl25b- expressing cells in a short period. As expected, few GFP positive cells were observed in the CMZ for such a short induction time wherein the clone partially overlaps with early retinal progenitor cells expressing rx2 and eventually produces late progenitors (Fig.S1E-E’). The proliferative capacity of *ccl25b* expressing RSCs was evaluated with BrdU labelling, showing that these cells are slowly proliferating as they did not incorporate BrdU when exposed for 1 day unlike the adjacent fast-cycling progenitor cells, which show BrdU incorporation (Fig.S1D-D’). *ccl25b* based iArCoSs were exclusively detected in the neuroretina. This validates *ccl25b* expressing cells as *bona fide* multipotent neuro-retinal stem cells residing in the peripheral tip of the CMZ.

**Figure 1:**
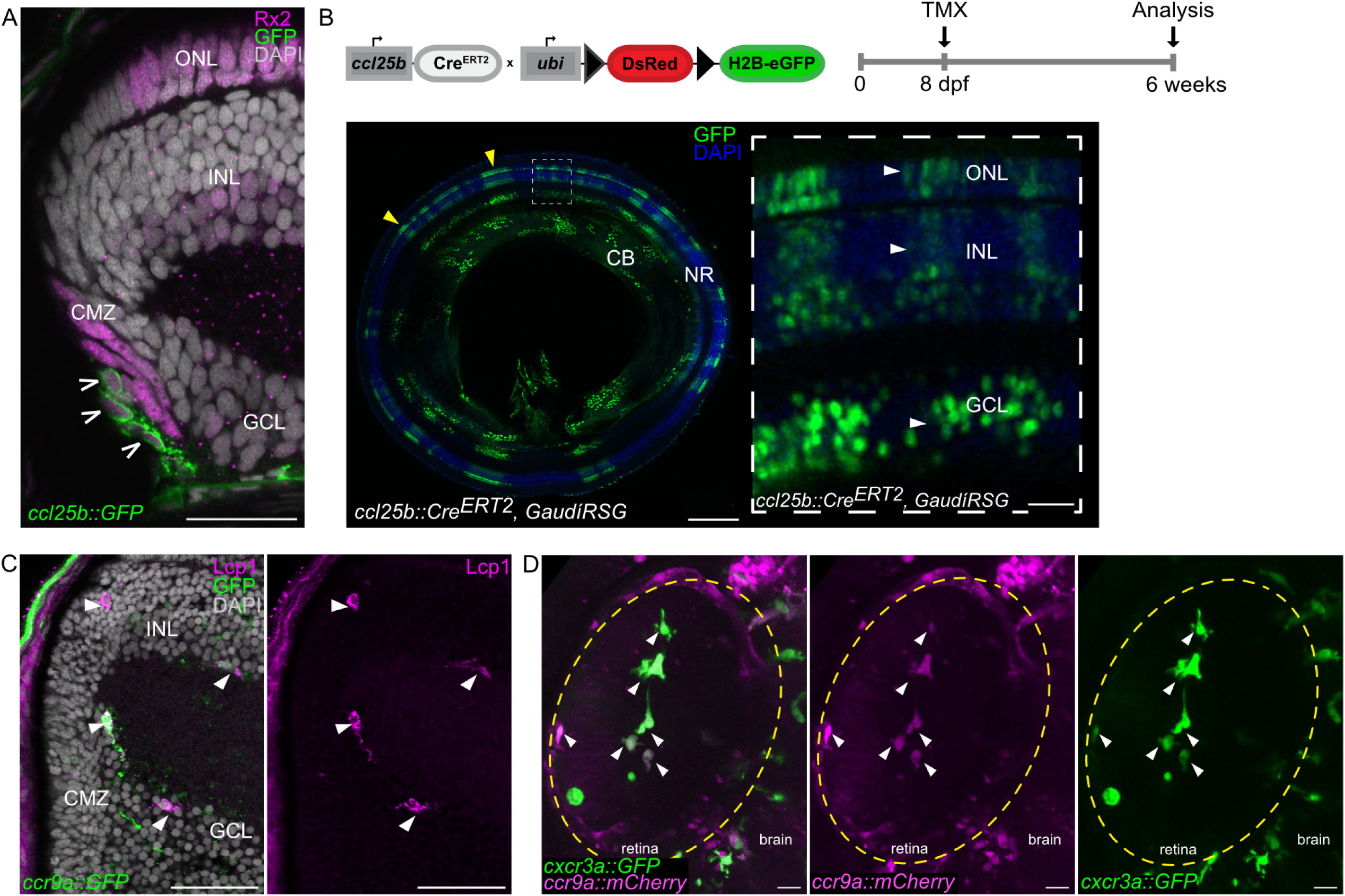
**Characterization of *ccl25b* as a retinal stem cell marker and *ccr9a* as marker for microglia in the retina.** (A) Transverse section of a retina derived from a *ccl25b::GFP* reporter hatchling. GFP- positive cells (open arrowheads) are located in the most peripheral cells of the CMZ and the Rx2-positive (magenta) domain, cell nuclei are stained with DAPI (grey). Scale bar: 20 µm. (B) Cre/loxP-mediated lineage tracing of *ccl25b*-positive cells: Genetic construct and induction scheme and wholemount *ccl25b::Cre^ERT2^ X GaudíRSG* retina, where recombination (red to green) was induced at hatching stage and analysed 6 weeks later. GFP-positive clones (iArCoS) (yellow arrowheads) can be detected in the neural retina (NR) and the ciliary body (CB). Scale bar: 200 µm. Zoomed in area of GFP-positive iArCoS (white dashed box) shows GFP+ cells in all neuroretinal layers (white arrowheads) in *ccl25b::Cre^ERT2^ X GaudíRSG* retinae. Scale bar: 20 µm. (C) Transverse section of a *ccr9a::GFP* hatchling retina. GFP positive cells are present in proximity to the CMZ region and in INL region. Lcp1 staining overlaps with *ccr9a::GFP* positive cells (white arrowheads). Cell nuclei are stained with DAPI (grey) Scale bar: 20 µm. (D) Co-expression of the macrophage reporter *cxcr3a::GFP* with *ccr9a::mCherry* in embryonic stage retinae (stage 28) as observed by *in vivo* confocal imaging. White arrowheads indicate co-expressing cells. Yellow dashed line indicate retina. Scale bar: 20 µm.

We complemented our analysis by probing for the presence and localization of cells expressing *ccr9a*, the cognate receptor of *ccl25b* in the retina of young fish at the hatching stage. We used an established transgenic line, *ccr9a::GFP* (Bajoghli et al., 2015) and detected GFP expressing cells around the CMZ and in the INL region of the retina (Fig. 1C). The endogenous expression of *ccr9a* in the retina was confirmed by HCR *in situ* staining (Fig.S1C).

*Ccr9a*-expressing cells were characterized as immune cells using immunostaining for lymphocyte cytosolic protein 1 (Lcp1) (Jones et al., 1998). Among all Lcp1-positive cells analyzed (n = 316 cells in 56 sections from 4 fish), 97% co-expressed the chemokine receptor ccr9a (Fig. 1C). *Ccr9a*-positive cells tagged with mCherry fluorophore were further validated as immune cells by the co-expression of *cxcr3a::GFP*, a well-established macrophage-specific marker (Aghaallaei et al., 2010) (Fig. 1D). Under homeostatic conditions, the only immune cells known to colonize the immune-privileged retina are primordial macrophages, which give rise to microglia (Herbomel et al., 1999, 2001).

### Retinal stem cells are phagocytosed by tissue resident microglia

To address if there are microglia- stem cell interaction in the retina, we first carried out a detailed analysis of the spatial arrangement of retinal stem cells (expressing *ccl25b*) and microglia expressing the cognate receptor *ccr9a or cxcr3a* in the retina of freshly hatched medaka larvae. We carefully analysed 3D reconstructions of retinae of double positive *ccl25b::H2B-RFP; ccr9a::GFP* reporter fish. While RSCs resided in a ring in the very periphery of the CMZ (open arrowheads in Fig. 2A, 3-D schematic), microglia formed a concentric double ring at the proximal edge of the CMZ close to inner plexiform layer (IPL) and outer plexiform layer (OPL). Each ring was composed of evenly spaced *ccr9a* or cxcr3a positive microglia contacting each other (Fig. 2A, 3D schematic, Movie S1).

**Figure 2:**
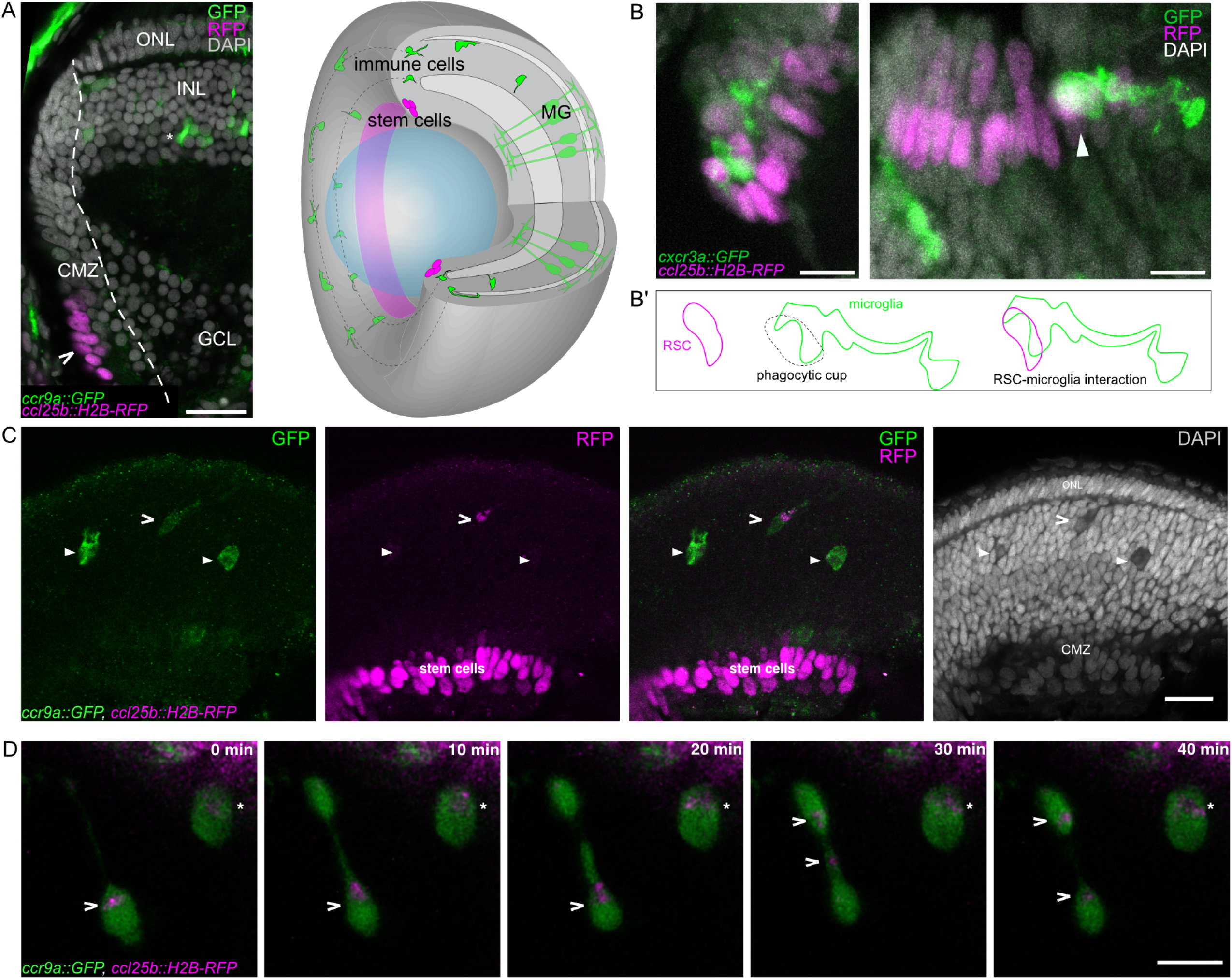
Ccl25b expressing RSC are phagocytosed by ccr9a+ microglia. (A) Transverse section of a *ccl25b::H2B-RFP*_*ccr9a::GFP* reporter retina of a hatchling. *Ccl25b*-driven H2B-RFP expression (open arrowheads) is located in the peripheral CMZ (dashed line). *Ccr9a*-driven GFP expression can be detected at the edge of the proximal end of the CMZ as well as in the INL of the retina (asterisk). Scale bar: 20 µm. 3D schematic representation of a retina expressing illustrating the distribution of *ccl25b* expressing RSCs (magenta) and *ccr9a* expressing microglia cells (green). (B) Retinal stem cell region showing interaction between H2B-RFP expressing *ccl25b+* (magenta) and GFP expressing *cxcr3a* positive cells (green), detected by antibody staining against RFP and GFP in wholemount retinae. Arrowhead indicates phagocytic cup in the process of forming. Scale bar: 10 µm. (B’) Schematic of the formation of a phagocytic cup by microglia to engulf RSC. (C) Wholemount staining showing the CMZ and retinal layers expressing H2B-RFP *in ccl25b* positive RSCs and GFP expressing in ccr9a+ microglia cells. White arrowheads depict *ccr9a* expressing cells enclosing H2B-RFP fractions (magenta) within phagosomes. Scale bar: 20 µm. (D) Time lapse of a *ccl25b::H2B-RFP* (magenta), *ccr9a::GFP* (green) retina depicting microglia transporting phagosomes within the CMZ (open arrowheads) and a static immune cell with a phagosome (asterisk). Scale bar: 10 µm.

An important feature of immune cells, particularly macrophages, is their capacity to phagocytose debris, damaged cells and foreign particles (Arandjelovic & Ravichandran, 2015; Nakayama, 2018). To address whether microglia phagocytose *ccl25b*-positive stem cells, we took advantage of the extended stability of red-fluorescent-protein (RFP)-tagged- H2B in the phagosomal compartment (Villani et al., 2019). We used a double transgenic line specifically labelling the nuclei of RSCs with histone-coupled RFP (*ccl25b::H2B-RFP*) and the microglia with GFP (*ccr9a::GFP*) (Fig. 2A). We found that GFP-labelled microglia migrate towards the RSC niche and get in direct contact with the *ccl25b* positive RSCs forming a phagocytic cup (Movie S3, Fig. 2B-B’). Microglia eventually contained RFP-positive phagosomes (Fig. 2C) and quantification of the phagocytosis of stem cells was carried out by counting the number of microglia containing stem cell derived RFP positive phagosomes. At hatching stage, we detected an active uptake by phagocytosis of stem cells in 61 % of the *ccr9a*-positive microglia (Fig. 3A’).

**Figure 3:**
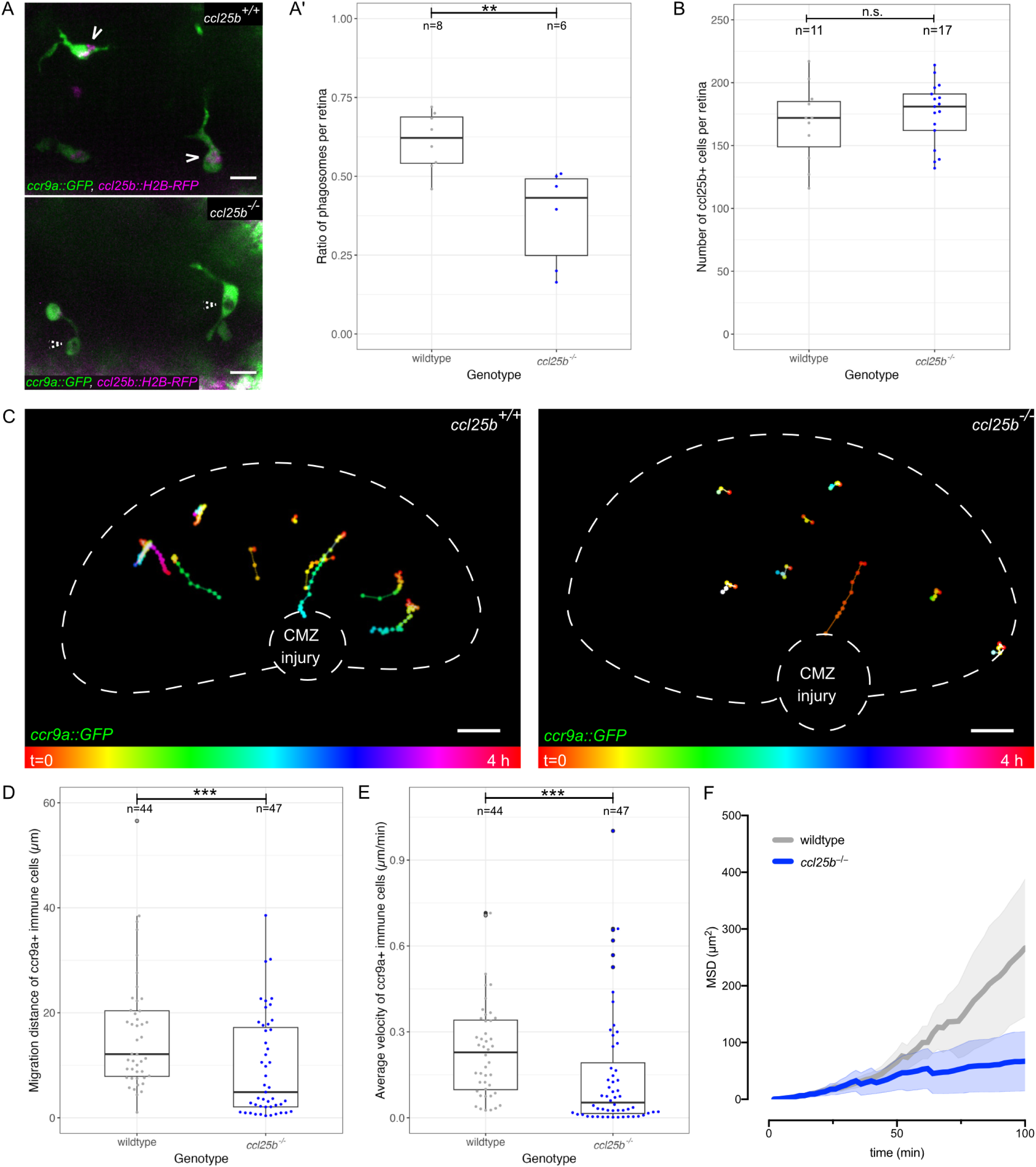
Ccl25b chemokine is necessary for activity of ccr9a positive immune cells in response to injury. (A) Representative images of phagosomes in wild type and *ccl25b^−/−^* retinae of *ccl25b:H2B- RFP* reporter fish. Scale bar: 10 µm. (A’) Comparison of percentage of *ccr9a*-positive immune cells with H2B-RFP-positive phagosomes in wildtype (n = 8) and *ccl25b^−/−^*(n = 6) retinae (**p = 0.004662; Mann-Whitney U test). (B) Quantification of *ccl25b+* retinal stem cells in wildtype (n=11) and *ccl25b^-/-^*(n=17) (p=0.4515; Mann-Whitney U test). (C) Representative tracks with temporal color-coding representing immune cell migration in *ccr9a::GFP* hatchling retinae in WT (n = 44 cells from 5 fish) and *ccl25b^−/−^* (n= 47 cells from 7 fish). Scale bar: 20 μm. (D) Quantification of *ccr9a* positive microglia migration distance in the retinae of *ccl25b^−/−^* vs wild type hatchlings (***p = 0.0008512; Mann Whitney U test). (E) Quantification of average migration velocity of microglia in the retinae of WT compared to *ccl25b^−/−^* hatchlings (***p = 0.0001817; Mann Whitney U test). (F) Analysis of mean square displacement (MSD) as a function of time from microglia cell tracking data. MSD of *ccl25b^−/−^*compared to MSD of microglia cells in WT retinae. Paler shading indicates standard error of means (SEM).

We investigated the dynamics of interactions between RSCs and microglia as well as their phagocytic activity using two-photon *in vivo* imaging at hatching stage. This analysis revealed microglia actively transporting stem cell derived, RFP-positive, phagosomes within the ciliary marginal zone (CMZ) (Fig. 2D, Movie S2). We observed microglia containing H2B-RFP-positive nuclei within their phagosomes, indicative for the direct uptake of stem cell nuclei (Fig. 2D, asterisk). We next addressed whether the phagocytosed stem cells represent cells being cleared through classical efferocytosis of apoptotic cells. Strikingly, TUNEL staining did not detect any apoptotic cells among RSCs. (Fig. S2C). This indicates that the detected prominent phagocytosis of RSCs is independent of apoptosis.

### Activity of microglia is dependent on *ccl25b* chemokine

The RSC specific expression of the chemokine ligand *ccl25b*, combined with our finding of *ccr9a* positive microglia-mediated stem cell phagocytosis, suggests that the ligand provides localized guidance cues that direct the activity of *ccr9a*-positive microglia.

To address the role of Ccl25b in the microglia mediated phagocytosis of RSCs we generated a *ccl25b^−/−^* mutant allele by targeted mutagenesis using CRISPR/Cas9. This allowed to test if the absence of Ccl25b affects the interaction between RSCs and immune cells. Targeted mutagenesis was designed to delete the majority of the *ccl25b* gene and the mutant transcript contained only 22 % of the wild type coding sequence. The mutated gene was lacking the CC motif crucial for folding and function of CC class chemokines (Thomas et al., 2015) (Fig. S2A).

The effect of loss of *ccl25b* functionality on phagocytosis, was assessed in *ccl25b^−/−^* mutants in the background of the *ccl25b::H2B-RFP*, *ccr9a::GFP* double reporter line. We examined the number of immune cells containing RSC-derived phagosomes in homeostatic conditions. We found that in wild type fish RSC-derived phagosomes were present in 61 % of *ccr9a*- positive cells (n = 201/331 cells in 8 retinae). In *ccl25b^−/−^* mutants the number of *ccr9a*- positive microglia containing RSC-derived H2B-RFP-positive phagosomes was reduced to 36 % (Fig. 3A’) (n = 113/317 cells in 6 retinae). Moreover, there was an increase in the number of *ccr9a*-positive microglia observed in the *ccl25b^−/−^* fish. Strikingly, the reduction in phagocytosis and the increase in the number of microglia in the absence of the Ccl25b chemokine did not affect the number of RSCs expressing *ccl25b::H2B-RFP* (Fig. S2B, Fig. 3B). The comparison of transverse sections of 3-week post hatch retinae between wildtype and *ccl25b^-/-^*, showed no gross morphological differences (Fig. S2D-D’).

Since chemokines are known to have a key function in acute injury responses, by attracting immune cells towards the site of inflammation(Sokol & Luster, 2015), we addressed whether Ccl25b is required to direct the migration of *ccr9a*-positive microglia within the retina to the stem cell domain upon injury. We used two-photon mediated RSC ablation to inflict a wound to the stem cell niche in the retina at hatching stage as has been described previously (Lust & Wittbrodt, 2018). We recorded the response of immune cells to the injury over four hours, with a temporal resolution of 2 min and analyzed their speed and directionality. In wild type retinae, *ccr9a*-positive microglia migrated rapidly to the injured area (Fig. 3C-F) while in *ccl25b^−/−^* retinae, the migratory behavior of *ccr9a*-positive microglia was significantly reduced and lacked directionality. In general, they displayed lower average migration velocity and travelled shorter total distances in the absence of Ccl25b (Fig. 3D-F).

These results indicate a crucial role of the Ccl25b-Ccr9a signalling axis in giving direction and specificity to microglia under inflammatory conditions and impacting on their ability to phagocytose stem cells without affecting the overall stem cell number under homeostatic condition.

### Absence of microglia affects the retinal stem cell numbers

Because phagocytosis was only modestly reduced in the absence of Ccl25b, we next asked whether microglia directly control RSC homeostasis. To eliminate microglia we targeted two transcription factors required for their differentiation, *spi1b* and *irf8*. While Spi1b is required for the development of both, lymphoid and myeloid lineages (Rhodes et al., 2005; Roh-Johnson et al., 2017; Scott et al., 1994), Irf8 is specifically involved during macrophage differentiation (Shiau et al., 2015).

Using CRISPR/Cas9-mediated mutagenesis, we generated a stable *spi1b^−/−^* mutant line and established and analyzed *irf8* crispants. The *spi1b* mutant carries a 6 bp insertion and 10 bp deletion in the 5^th^ exon of *spi1b*, resulting in a premature stop codon (Fig. S3A-A’) truncating the C-terminus of the Spi1b protein. Loss of Spi1b resulted in the complete absence of macrophages as validated by Sudan black and Neutral red staining. In *spi1b^−/−^* homozygous mutants, Sudan Black stained blue-black cells were absent from the tail fin (Fig. S3B-B’). Neutral red staining which specifically labels lysosome-rich cells such as macrophages was observed in the tail fin of wildtype but not the *spi1b^-/-^* fish (Fig. S3C-C’). Additionally, in situ hybridization with a macrophage specific marker *cxcr3a*, labelled macrophages in the brain, gills, intestine and skin of control wildtype hatchlings but gave no signal in *spi1b^-/-^*fish (Fig. S3D-D’). These results validate the complete absence of macrophages upon loss of Spi1b function. The loss of *spi1b* is lethal at juvenile stages (3 weeks post hatching) and the overall morphology of the retina is disrupted by this stage (Fig. S3F).

To determine whether microglia regulate the RSC niche in the CMZ, we introduced the *ccl25b::H2B-RFP* reporter into the *spi1b^+/−^* mutant background. Loss of microglia resulted in a pronounced expansion of the RSC population already at the hatching stage, reflected by an increased width of the ccl25b-positive CMZ (Fig. 4A-A’). Quantification across the entire CMZ revealed a 34.3% increase in RSC number in the absence of microglia (*spi1b^−/−^*: n = 13 retinae; wild-type siblings: n = 51 retinae) (Fig. 4A’’).

**Figure 4:**
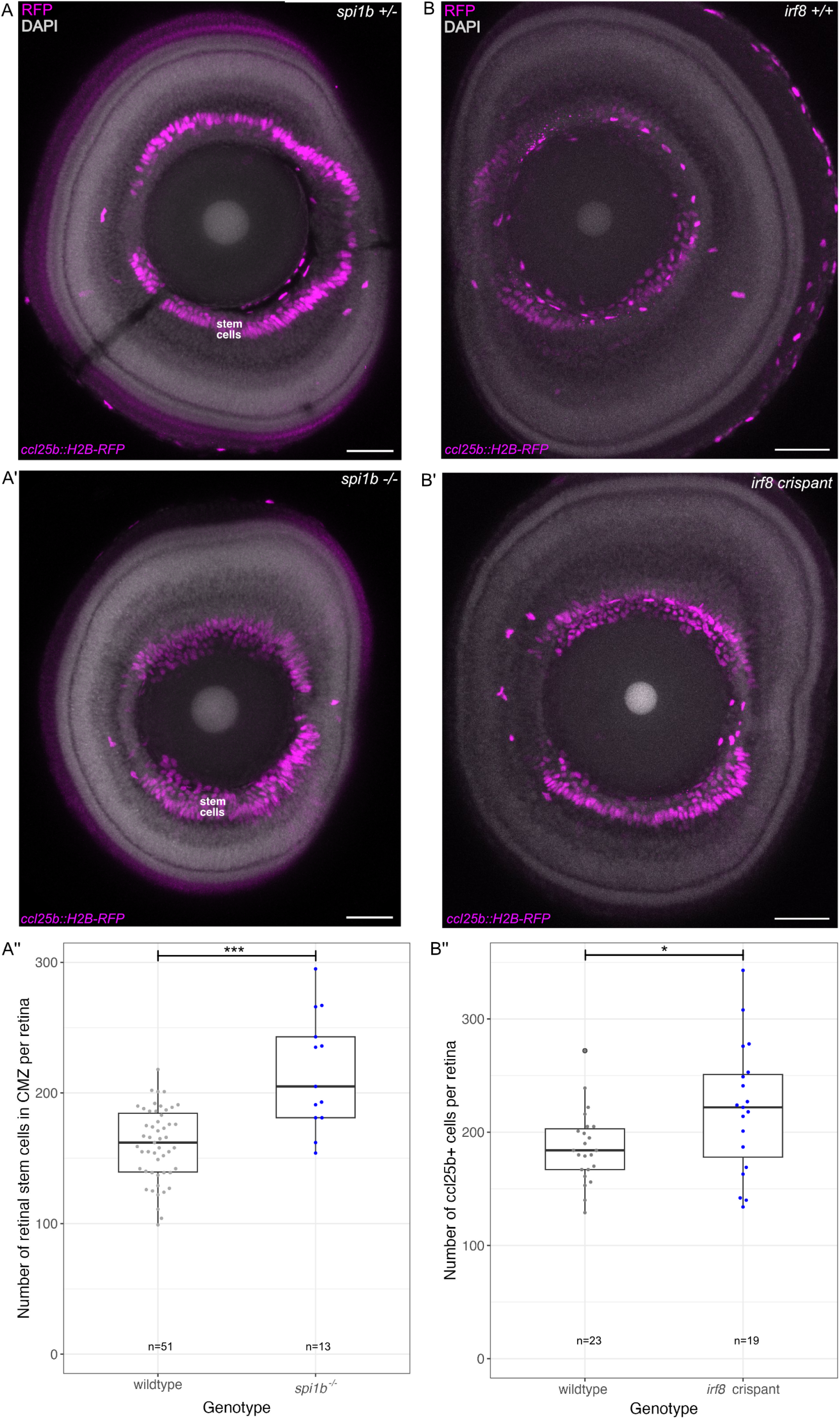
Ccl25b retinal stem cells are affected in absence of microglia. (A-A’) Sagittal view on retinae of *ccl25b::H2B-RFP* reporter hatchlings in WT and *spi1b^-/-^* background. H2B-RFP was detected by antibody staining of whole retina (magenta). Scale bar: 50 µm. (A’’) Comparison of *ccl25b* expressing retinal stem cells in wildtype hatchings (n=51 fish) and *spi1b^-/-^* hatchlings (n=13 fish) (***p = 0.0000908; Mann-Whitney U test). (B-B’) Sagittal view on retinae of *ccl25b::H2B-RFP* reporter hatchlings in WT and *irf8* crispant. H2B-RFP was detected by antibody staining of whole retina (magenta). Scale bar: 50 µm. (B’’) Comparison of *ccl25b* expressing retinal stem cells in wildtype hatchings (n=16 fish, 23 retinae) and *irf8* crispant hatchlings (n=10 fish, 19 retinae) (*p = 0.02878; Mann-Whitney U test).

To distinguish microglia specific effects from the potentially broader immune defects caused by the loss of *spi1b*, we independently depleted microglia by CRISPR/Cas9 mediated targeting of the microglia-specific transcription factor *irf8*. *Irf8* crispants phenocopied the *spi1b* mutants and displayed an 18.1% increase in the number of RSCs (*irf8* crispant: n = 19 retinae; wild-type siblings: n = 23 retinae) (Fig. 4B’’). This demonstrates that the expansion of the stem cell pool is a direct consequence of the loss of microglia.

The enlarged RSC compartment persisted throughout larval development until the death of the *spi1b^−/−^* mutants, presented here in an exemplary section of the CMZ in a young fish 5 days post hatching (Fig. 5A-A’). At the tissue level, mutant larvae displayed smaller, frequently asymmetric retinae, while the overall body length was only mildly affected (Fig. 5B, Fig. S3E).

**Figure 5:**
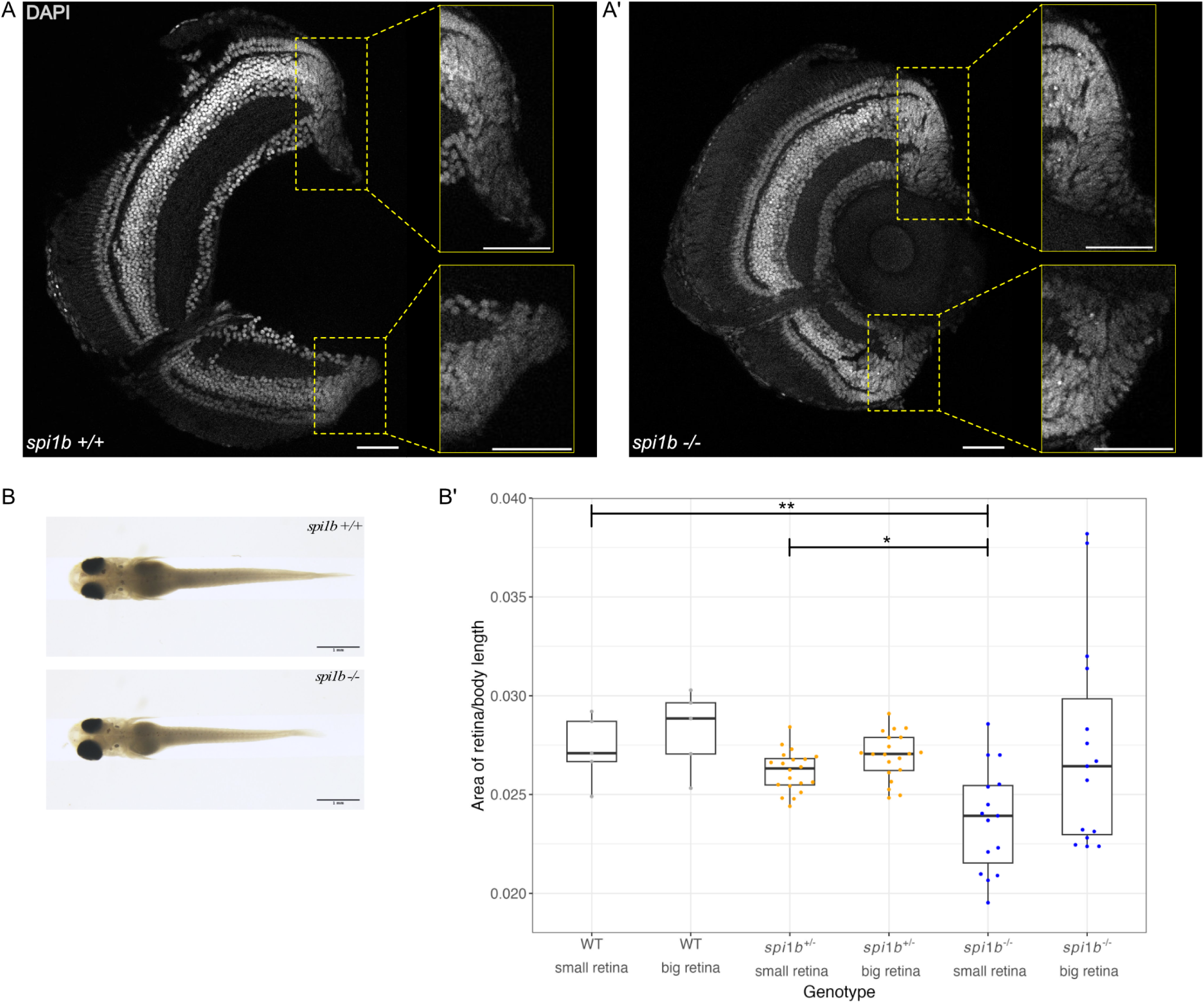
Size of CMZ and retina are affected in the absence of microglia. (A-A’) Transverse section of wildtype and *spi1b^-/-^* retina stained with DAPI (grey) at 5 days post hatch. CMZ region is highlighted and zoomed in (yellow box) showing nuclei organization. Scale bar: 50 µm. (B) Dorsal view of wildtype and *spi1b^-/-^* hatchlings to visualize differences in the size of retina. Scale bar is 1 mm. (B’) Quantification of size of the retina by area normalized with the length of the fish to see any differences between wildtype (n=5), spi1b^+/-^ (n=20) and spi1b^-/-^ (n=15). The retina sizes are grouped into small and big retina for every hatchling. (**p = 0.008900154 for WT small retina vs. spi1b^-/-^ small retina, *p = 0.013824911 for spi1b^+/-^ small retina vs. spi1b^-/-^ small retina; Dunn test with Holm- Bonferroni correction).

Thus, despite the expansion of the retinal stem cell pool, retinal growth is impaired in the absence of microglia indicating that microglial uptake regulate retinal stem cell numbers to ensure proper tissue architecture under homeostatic growth conditions.

## Discussion

Our findings identify microglia as key regulators of retinal stem cell (RSC) homeostasis in the medaka retina. The life-long activity of the RSC niche, marked by the expression of *ccl25b*, requires continuous extrinsic regulation by microglia through an apoptosis independent mechanism. Loss of microglial surveillance in *spi1b^−/−^* mutants and irf8 crispants, results in pronounced expansion of the RSC population and consequently an enlarged stem cell niche (CMZ). Surprisingly, this increase in stem cell number is accompanied by a reduction in overall growth during post-embryonic development. These opposing phenotypes suggest that stem cell abundance alone is not sufficient to sustain coordinated tissue growth. Instead, excessive RSC density may impair the balance between self-renewal and differentiation ultimately compromising retinal growth. We therefore hypothesize that microglia maintain an optimal RSC population size that preserves the functional output of the stem cell niche. Consistent with this notion, the uneven expansion of RSCs around the mutant CMZ leads to anisotropic retinal growth, demonstrating that local regulation of stem cell abundance directly translates into altered tissue morphogenesis within the same individual.

Although *spi1b* broadly controls the differentiation of myeloid and lymphoid lineages, whereas *irf8* is more specifically required for macrophage and microglial differentiation, disruption of either gene resulted in a comparable expansion of the RSC population. Since the molecular mechanisms governing microglial differentiation are heterogenous and can vary between different developmental stages, we focused a large part of the analysis on our established stable *spi1b* mutant line. This strategy also avoided reliance on a single microglial differentiation pathway that may affect only a subset of of microglia and therefore provides greater confidence that the observed phenotype reflects the consequences of robust and sustained microglial depletion rather than defects in a specific microglial subpopulation.

While the Ccl25-Ccr9 signalling pathway had previously been associated with antigen presentation in the intestine and T cell development in the thymus (Aghaallaei et al., 2022; Baubak Bajoghli, 2009), our observations demonstrate an additional function. We show that microglia, an innate immune cell that has not been previously linked to this pathway, express the *ccr9a* receptor and is involved in direct stem cell interaction. This highlights the tissue-specific role of Ccl25-Ccr9 signalling and its ability to influence both myeloid and lymphoid cell types differently.

Loss of Ccl25b specifically impairs microglial migration towards the site of injury in the CMZ and disrupts homeostatic phagocytosis of retinal stem cells. Although chemokines are known to facilitate immune cell homing during development (Wurbel et al., 2007; Wurbel et al., 2001), the absence of *ccl25b* in mutant retinae led to an increase in the number of *ccr9a* microglia present at the interface between CMZ and central retina, indicating that the presence of Ccl25b chemokine ensures proper number of *ccr9a* microglia in the retina. While microglia actively express the *ccl25b* receptor *ccr9a*, and an inflammatory response after injury is clearly affected without Ccl25b, this pathway is not essential for regulating stem cell numbers. Even though there is an increase in retinal microglia, the reduction in phagocytic capacity argues for the existence of an alternative mechanism of stem cell-microglia interaction for homeostatic stem cell maintenance.

While the knowledge about interaction between immune and stem cells has been rather limited, a similar regulatory mechanism by immune cells has recently been described for the hematopoietic stem cell niche (HSC) in zebrafish (Wattrus et al., 2022). The underlying mechanisms functioning in neuronal stem cells and microglia interaction in the medaka retina remain to be explored.

The teleost retina exhibits continuous, highly organized growth dependent on tightly regulated stem cell activity, and we demonstrate that microglia maintain this balance by phagocytosing RSCs. A fundamental question is whether this regulation serves as a quality control mechanism where specific cells display markers targeting them for phagocytosis, or if the process is stochastic, based on physical constraints within the CMZ similar to other developmental processes where cell elimination occurs through mechanical or signaling mechanisms (Huels et al., 2018; Levayer et al., 2016). This newly discovered role of the immune system opens exciting possibilities for understanding its involvement in the transition to and maintenance of multicellularity during development, providing an excellent foundation for exploring key mechanisms in stem cell niche regulation.

## Materials and Methods

### Fish husbandry and transgenic lines

Medaka (*Oryzias latipes*) were kept as closed stocks at Heidelberg University. All experimental procedures and husbandry were performed in accordance with the German animal welfare law and approved by the local government (Tierschutzgesetz §11, Abs. 1, Nr. 1, husbandry permit AZ 35–9185.64/BH and line generation permit AZ 35–9185.81/G-145-15). Fish were maintained in a constant recirculating system at 28°C on a 14 h light/10 h dark cycle. The following previously described stocks and transgenic lines were used for this study: wild-type Cab, *rx2::H2B-RFP* (Reinhardt et al., 2015), *ccr9a::GFP* (Bajoghli et al., 2015), *ccl25b::GFP, ccr9a:mCherry, ccl25b::H2B-RFP* (Aghaallaei et al., 2022), GaudíRSG (Centanin et al., 2014) and cxcr3a::GFP (Aghaallaei et al., 2010).

The following lines were generated for this study: *ccl25b*^-/-^, *ccl25b::Cre^ERT2^*, *spi1b^-/-^*. Transgenic lines were created by microinjection with Meganuclease (I-SceI) in medaka embryos at the one-cell stage, as previously described (Thermes et al., 2002).

### CRISPR/Cas9 mediated mutation of the *ccl25b*, *spi1b* and *irf8* locus

For generating truncated variants of *ccl25b* and *spi1b* we used CRISPR/Cas9 mediated mutation. The medaka *ccl25b* gene (Gene ID: ENSORLG00000028881, Exon ID: ENSORLE00000270681) comprises six exons. We selected two sgRNAs targeting the CC motif in Exon 4. The following target sites were used (PAM in brackets): GCGCCAATCCAGAGGACCCG(TGG) and CTTGGCTACGTGAGAGAACT(CGG). For mutation of Spi1b we used one sgRNA targeting exon 5 of the *spi1b* gene: GAGGCCCTCGCACACCGCTGGG (Gene ID: ENSORLG00000006090, Exon ID: ENSORLE00000068479. For creating *irf8* crispants (Gene ID: ENSORLG00000018216.2, Exon ID: id73007.1) a sgRNA targeting exon 2 of the *irf8* gene was used. All sgRNAs were designed using the CCTop prediction tool (Stemmer et al., 2015). Cloning of sgRNA templates and in vitro transcription was performed as described previously. The pCS2+Cas9 plasmid was linearized using NotI and the mRNA was transcribed *in vitro* using the mMessage mMachine SP6 kit (ThermoFisher Scientific, AM1340). The CRISPR generated edits were mapped and analysed using geneious software version 2019.2.3 (www.geneious.com).

One-cell stage medaka were injected with 150 ng/µl of Cas9 mRNA and 15 ng/µl of the sgRNAs. Injected embryos were maintained at 28°C in embryo rearing medium (ERM, 17 mM NaCl, 40 mM KCl, 0.27 mM CaCl_2_, 0.66 mM MgSO_4_, 17 mM Hepes). One day post injection (dpi) embryos were screened for survival.

### Genotyping and mutant selection

For genotyping either injected F0 embryos or fin clips of juvenile or adult fish were lysed in DNA extraction buffer (0.4 M Tris/HCl pH 8.0, 0.15 M NaCl, 0.1 % SDS, 5 mM EDTA pH 8.0, 1 mg/ml proteinase K) at 60°C overnight. Proteinase K was inactivated at 95°C for 10 min and the solution was diluted 1:2 with H_2_O.

The PCR amplification was performed with Q5 polymerase (NEB) according to the manufacturers protocol using 30 PCR cycles. The resulting PCR product was analysed on a 1% agarose gel. F0 animals were raised to adulthood and outcrossed to wild type Cab fish. F1 fish were then raised to adulthood and finclipped and genotyped. Mutants carrying the same allele were incrossed to obtain homozygous mutants.

To identify mutations in the locus, fin clip DNA of F1 fish was analysed as described above. Genotyping of ccl25b mutants revealed a 333 bp deletion in the ccl25b locus, completely eliminating exon four including the CC motif along with two-thirds of exon five (Fig. S2). Transcript analysis of wild type and *ccl25b^−/−^*fish showed that the resulting truncated transcript contained only 22% of the wild type coding sequence. We selected *spi1b^−/−^* mutants carry a 6bp insertion and 10bp deletion in exon 5 which leads to a frame shift and premature stop codon. In case of *irf8* crispants, fin clip DNA of F0 hatchlings was used and sent for sequencing

### Tamoxifen induction of Cre/lox recombination in ccl25b expressing cells

For Cre^ERT2^ induced recombination of *ccl25b* expressing cells, *ccl25b::Cre^ERT2^* X *GaudiRSG* (Centanin et al., 2014) hatchlings (0 day post hatch) were treated with 5 µM tamoxifen solution (Sigma-Aldrich) in 1x ERM overnight. Fish were then grown for 3 days or six weeks before they were sacrificed for whole mount immunostaining of retinae as described below.

### Immunohistochemistry on whole-mount retinae

*ccl25b::Cre^ERT2^ X GaudiRSG* fish grown for 6 weeks after induction of recombination were euthanized with 20x Tricaine (Stock: 4 g/l triciane; 10g/l Na2HPO4·2 H2O; pH 7-7.5), fixed overnight in 4 % PFA and treated and stained as described in(Perez Saturnino et al., 2018). To detect GFP after Cre-induced recombination in *ccl25b* expressing cells we used chicken anti-eGFP (1:500, Life Technologies, A10262) and donkey anti-chicken Alexa Flour 488 (1:750, Jackson, 703-485-155) and DAPI (2 mg/ml) (1:500). Retinae of *cxcr3a::GFP;ccl25b::H2B-RFP* reporter hatchlings were treated and stained as in (Perez Saturnino et al., 2018) except bleaching time was reduced to 1h.

For the detection of GFP, we used chicken anti-eGFP (1:500, Life Technologies, A10262) and donkey anti-chicken Alexa Flour 488 (1:750, Jackson, 703-485-155). For detection of RFP, we used rabbit anti-RFP (1:500, Clontech 632496) as primary antibody and goat anti-rabbit DyLight549 (1:750, Jackson, 112-505-144) or goat anti-rabbit Alexa Fluor 647 (1:250, Thermo LifeTech, A21245) as secondary antibody. To visualize nuclei organization, retinae of *spi1b^+/-^*and *spi1b^-/-^* hatchlings were stained for DAPI (1:500) at 5 days post hatch.

### BrdU incorporation assay

To follow cell division in *ccl25b* positive cells we incubated *ccl25b::GFP* reporter hatchling in 2.5 mM BrdU diluted in 1x Embryo Rearing Medium (ERM) for 1 day at 28°C. After removal of BrdU fish were euthanized, fixed and treated for cryo-sectioning.

### Immunohistochemistry on cryosections

To obtain cellular resolution of the CMZ and surrounding cell layers, immunohistochemistry was performed on hatchlings as described previously(Lust & Wittbrodt, 2018). For the detection of GFP in *ccr9a:GFP* we used chicken anti-eGFP (Life Technologies, A10262) and donkey anti-chicken Alexa Flour 488 (Jackson, 703-485-155). For the detection of BrdU cryosections were fixed in 4% PFA (paraformaldehyde; pH 7) for 30 mins after the incubation with secondary antibody and DAPI. After fixation sections were washed 3x for 10 mins with 1x PTW. The sections were then treated with 2N HCl, 0.5% Triton X in 1x PBS for 60 min at 37°C for antigen retrieval, washed again 3x for 10 mins with 1x PTW and then treated with Borax/PTW for 15 mins for pH recovery. After washing 3x for 10 mins sections were blocked with 10% NGS in 1x PTW for 2 hours, then washed 2x for 5 mins with 1x PTW before applying the primary antibody (1:100 or 1:200 BrdU (Abcam, ab6326)) in 1% NGS) overnight at 4°C. The next day the sections were washed 6x for 10 mins with 1x PTW before secondary antibody was applied (1:750 anti-rat DyLight488 (Jackson, 112-485-143) in 1% NGS) for 2 hours at 37°C and sections were washed again with 1x PTW before mounting as described in (Lust & Wittbrodt, 2018).

### In situ hybridization chain reaction (HCR)

HCR probe sets for *ccl25b*, *rx2* and *ccr9a* were designed against unique mRNA sequences, based on the Japanese medaka HdrR ASM223467v1 transcriptome. Probes (50pmol) were ordered as IDT Oligo pools. Alexa-488 and Alexa-594 conjugated hairpins were purchased from Molecular instruments together with the buffers. CUBIC-based clearing method was used to improve probe penetration (Susaki & Ueda, 2016 and http://www.cubic.riken.jp). Fixed and washed medaka hatchlings were incubated in 50% CUBIC-L/R1a (mixed 1:1 and then diluted to 50% in dH2O) for 30 mins at room temperature while shaking and moved to a 100% CUBIC-L/R1a solution for another 30 min incubation at room temperature on a shaker. Afterwards the embryos were washed 4 times 15 minutes with 1x PBS. HCR was performed according to the Molecular Instruments instructions. Probe concentration was increased to 4 pmol. Embryos were incubated with DAPI (1:500 in 5X SSCT) for 10 minutes at room temperature, transferred to a MatTek glass-bottom dish and equilibrated in CUBIC refractive index matching solution CUBIC R+(N) until transparent.

Imaging was performed using the 63x objective on Leica SP8 confocal microscope. Image processing was done using Fiji. Bright outliers (of 1 pixel radius) were removed to denoise the image. Plugin “Subtract background” was used to correct background (rolling ball diameter = 75 pixels). Maximum projection of 6-10 z-slices (1µm z-step).

### TUNEL staining

To measure the amount of apoptotic cells, we made use of In situ cell death detection kit,TMR red (Roche, 12156792910). To stain cryosections, 50 µl TUNEL reaction mix was added per slide and incubated at 37°C for 1 hr. Slides were washed twice with 1X PTW and mounted in 60% glycerol. For whole mount staining, retinae was permeabilized using 0.1% Triton X-100, 0.1% sodium citrate for 30 min at RT. Samples were washed twice with 1X PTW and incubated with the TUNEL reaction mix at 37°C for 1 hr in the dark. After washing twice with 1X PTW, they were mounted for imaging.

### In vivo imaging and laser induced injury

For in vivo imaging, transgenic Cab embryos were kept in 5x 1-phenyl-2-thiourea (PTU (0.0165% m/v), Sigma) in 1x ERM from 1 dpf until imaging to block pigmentation. For embryonic stage imaging, stage 34 embryos were hatched using labmade hatching enzyme. Live hatchlings were anaesthetized in 1x Tricaine diluted in 1x ERM and mounted in glass bottomed Petri dishes (MaTek) in 1 % low melting point agarose.

In vivo imaging and laser ablations were performed on a Leica TCS SP8 and SP5 microscope equipped with a Spectra Physics Mai Tai® HP DeepSee Ti:Sapphire laser, which has a tuneable wavelength range from 690-1040 nm, and Leica Hybrid Detectors. The retinal injury was induced using a 14x zoom in combination with the high-energy 2-photon laser, precisely tuned to 880 nm. Subsequent follow-up imaging for GFP detection was conducted using the identical laser wavelength of 880 nm and a 40x long distance objective. Imaging of double reporter fish was performed on a Leica DIVE microscope.

### Sudan black staining

For visualizing macrophages in medaka hatchlings we used Sudan black staining as described in(Aghaallaei et al., 2022). Medaka embryos were fixed in 4% PFA overnight, washed with PTW and incubated in Sudan black staining reagent (Sigma Aldrich 380B-1KT) for 1 hour in the dark. After staining, embryos were washed 3x with 70% EtOH and 3x with PTW. Pigments were bleached using 3% H2O2 + 0.5 % KOH for 15 min and embryos were washed again 3x with PTW before they were counterstained with Hematoxylin staining solution for 1 hour. Whole hatchling stained with Sudan black were captured using a Nikon SMZ18 Stereomicroscope fitted with a Nikon DS-Ri1 camera.

### Neutral red staining

Medaka embryos from *spi1b* mutant line were grown until hatchling stage (1 dph) in 1x ERM at 28°C. Hatchlings were stained live with 2.5 µg/ml of Neutral red solution (Sigma Aldrich 72210) prepared in 1x ERM for 6 h at 28°C. After staining, hatchlings were rinsed with 1x ERM and anaesthesized using 2X Tricaine solution. Hatchlings were mounted in 3% methylcellulose and imaged using Nikon Eclipse 80i microscope. This staining protocol was established as described in (Wei et al., 2024).

### In situ hybridization

For visualizing macrophages in medaka hatchlings whole mount *in situ* hybridization was performed using digoxygenin labelled *cxcr3a* probe transcribed from the cDNA of the second exon of the gene (ENSORLG00000013459.2). Medaka hatchlings from *spi1b* mutant line were fixed in 4% PFA overnight, washed with PTW and permeabilized with Proteinase K for 10 min. Whole mount *in situ* hybridization using NBT/BCIP solution was carried out as described previously (Loosli et al., 1998). After staining, hatchlings were washed 3x with PTW and mounted on microscopic slides with 60% glycerol in PBS. Images were acquired using Nikon Eclipse 80i microscope.

### Immunohistochemistry imaging

All cryosections and whole mount samples were imaged using an inverted confocal laser scanning microscope Leica TCS SP8. The microscope was equipped with ACS APO objective lenses (10x/0.3 dry, 20x/0.75 multi-immersion, 63x/1.3 glycerol) and featured laser lines at 405 nm, 488 nm, 532 nm, and 638 nm. The system was configured with two PMT detectors. For whole mount sample imaging, retinae were mounted in MatTek dishes using 1% low melting agarose in 1x PTW or clearing solution ((Zhu et al., 2019)(20% (wt/vol) urea (Sigma–Aldrich, Cat^#^: V900119), 30% (wt/vol) D sorbitol (Sigma–Aldrich, Cat^#^: V900390), 5% (wt/vol) glycerol (Sigma–Aldrich, Cat^#^: V900122) dissolved in DMSO (Sigma Aldrich, Cat#: V900090))., with the Ciliary Marginal Zone (CMZ) positioned touching the bottom of the dish. After the agarose solidified, the dish was covered with 1x PTW. No solution was added on top in case of clearing solution.

### Image acquisition of hatchlings

To measure the size of the retina and length of the *spi1b* wildtype and mutant hatchlings, samples were fixed in 4% PFA in PTW and mounted in 1.5% agarose plates with customized wells. Images were acquired using Nikon SMZ18 microscope with DS-Ri1 camera.

### Image processing and statistical analysis

Images were processed using the Fiji image processing software. Statistical analysis and graphical representation of the data were performed using Rstudio and Prism software package (GraphPad)(*RStudio: Integrated Development Environment for R.*, 2023). Tracking of immune cells was carried out using MtrackJ plugin in Fiji. Average velocity, total migration distance and mean square displacement (MSD) as a function of time were calculated for each immune cell. The MSD at time point t was calculated as the squared distance between immune cell position at time point t and at t = 0. Box plots show median, 25^th^ and 75^th^ percentiles. Cell counting was done using cell counter plugin in Fiji. Unpaired t-tests and Mann Whitney test were performed to determine the statistical significances. The P value P < 0.05 was considered significant and P values are given in the figure legends. Sample size (n) is mentioned in figure legends. The experimental groups were allocated randomly, and no blinding was done during allocation. The figures in this study were created using the software Affinity Designer 2, version 2.5.5 (2024), Serif.

## Acknowledgments and funding

We thank the Wittbrodt lab for constructive discussions on the project, Lazaro Centanin, Carmen Gloria Feijoo, Tinatini Tavhelidse-Suck, Thomas Thumberger, Erika Tsingos and Lucie Zilova for various contributions and critical reading of the manuscript. We are grateful to Leica Microsystems Germany, in particular K.-H. Körtje for providing access to and assistance with the Leica DIVE imaging setup. We are grateful to Tanja Kellner and Beate Wittbrodt for expert technical assistance. Pimpan Sujariyakul for work on establishing the *spi1b* mutant line and Nithyapriya Kumar for studying the embryonic and post-embryonic phagocytosis behavior. Jana Fuß for providing the data on Ccl25b expression on Müller glia cells. We thank Erik Leist and Marzena Majewski and Antonino Saraceno for fish husbandry. K.L., C.B. and R.A. were members of HBIGS, the Heidelberg Graduate School for Life Sciences. K.L. was supported by a LGFG Fellowship by the Ministerium fu r Wissenschaft, Forschung und Kunst Baden-Wu rttemberg. C.B. was supported by a PhD fellowship of the Studienstiftung des deutschen Volkes. O.S. was supported by an HBIGS MSc fellowship. This work was supported by the European Research Council (GA 294354-ManISteC and synergy grant 810172/IndiGene to J.W.) and the Deutsche Forschungsgemeinschaft (SFB 873 TP A3 to J.W.).

## Author contributions

R.A., K.L., C.B., J.B and J.W. conceived the study and designed the experiments and J.W. supervised the work. R.A., K.L., C.B., J.B., N.F., F.E. and O.S. performed, discussed and refined the experiments in constant discussion with J.W.. B.B. and N.A. contributed transgenic fish lines and conceptual input. R.A., K.L., C.B., J.B., E.HdC. and J.W. wrote the manuscript with feedback from all authors.

## Data and materials availability

All data are available in the main text or the supplementary materials.

