## Supplementary figures for "Immune regulation of neuronal stem cells in the medaka retina"

**Supplementary Materials:**

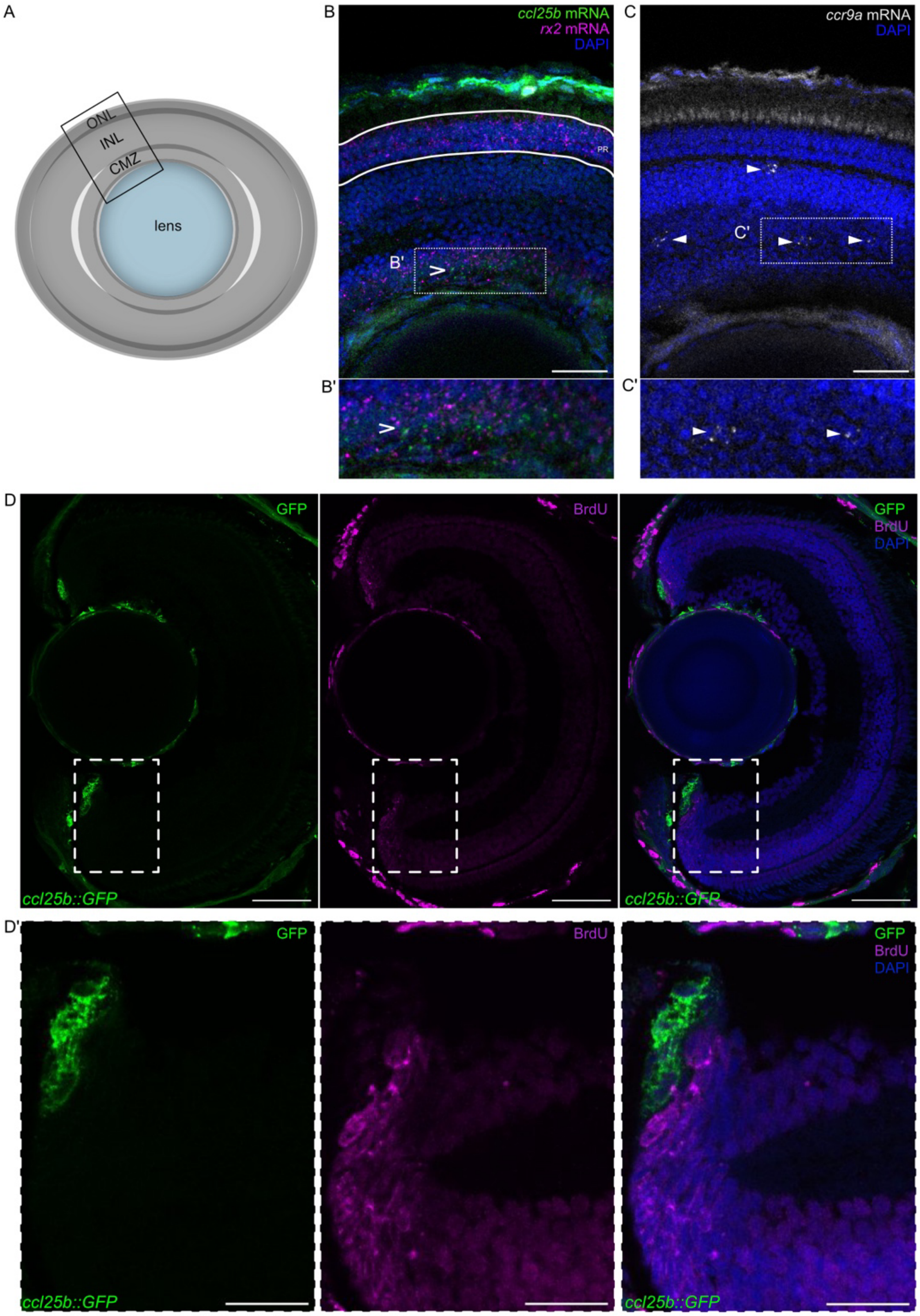

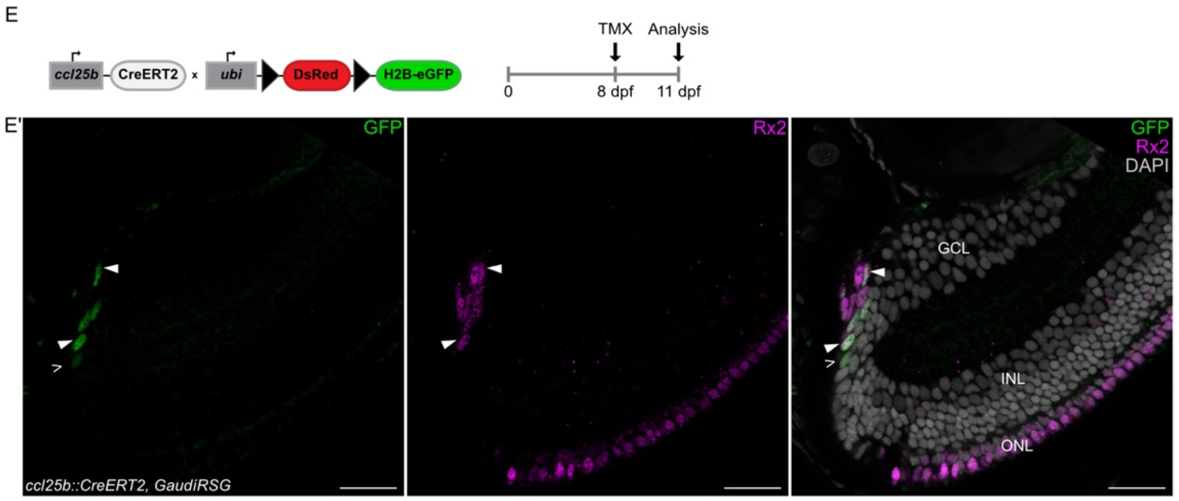


**Figure S1: ccl25b and ccr9a endogenous expression in the retina and proliferative behaviour of ccl25b+ cells**

(A) Schematic diagram of a whole retina showing the different regions: the CMZ (stem and progenitor cell region) and inner and outer nuclear layers (INL and ONL) (differentiated regions). Box shows the area of the retina represented in B and C. (B) Hybridisation chain reaction (HCR) *in situ* staining of *ccl25b* and *rx2* mRNA to detect their endogenous expression in differentiated retinal layers and the CMZ region (B'). Scale bar: 20 µm. (C) HCR *in situ* staining of *ccr9a* mRNA in the retinal layers and specifically in the INL (white box, C'). Scale bar: 20µm. (D) Transverse section of a *ccl25b::GFP* hatchling retina stained for BrdU incorporation after a short 24 h BrdU pulse. BrdU positive cells in relation to *ccl25b* expressing RSC at the periphery of CMZ. Scale bar: 50 µm. (D') Marked area of CMZ as shown in D (white box). Scale bar: 20 µm. (E-E') Earliest induction of *ccl25b::Cre* expressing cells (3 days post induction). Recombined cells express GFP and co-localise with rx2 positive cells (white filled arrows) at the beginning and eventually do not co-localise with rx2 (white line arrow) as the recombined stripe grows. Scale bar: 20µm.

**Movie S1: Distribution of microglia throughout the retina.** Microglia expressing *cxcr3a::GFP* are visualized using immunostaining in a post-embryonic whole mount retina (0 days post hatch). They are mainly positioned at the proximal edge of the CMZ near inner plexiform layer (IPL) and outer plexiform layer (OPL) as two concentric rings. In addition, they are also seen in other regions since they are migratory cells. White circle: lens, yellow circle: first ring of microglia near IPL, magenta circle: second ring of microglia near OPL. Scale bar is 50µm.

**Movie S2: *Ccr9a*-positive immune cells with H2B-RFP positive phagosomes derived from ccl25b expressing cells.**

*In vivo* imaging of a *ccl25b::H2B-RFP* (magenta), *ccr9a::GFP* (green) retina depicting mobile and static microglia containing H2B-RFP positive phagosomes within the CMZ and a static immune cell with a phagosome. Scale bar: 10 µm.

**Movie S3: Microglia interacts with retinal stem cells expressing *ccl25b::h2BRFP*.** In vivo imaging of an embryonic stage retina where microglia labelled with *cxcr3a::GFP* (green) interacts with *ccl25b* expressing retinal stem cells (magenta) by migrating in and out of the CMZ (interaction point at 01h 02min). Time interval is every 5 min 24 sec. Scale bar is 20 µm.

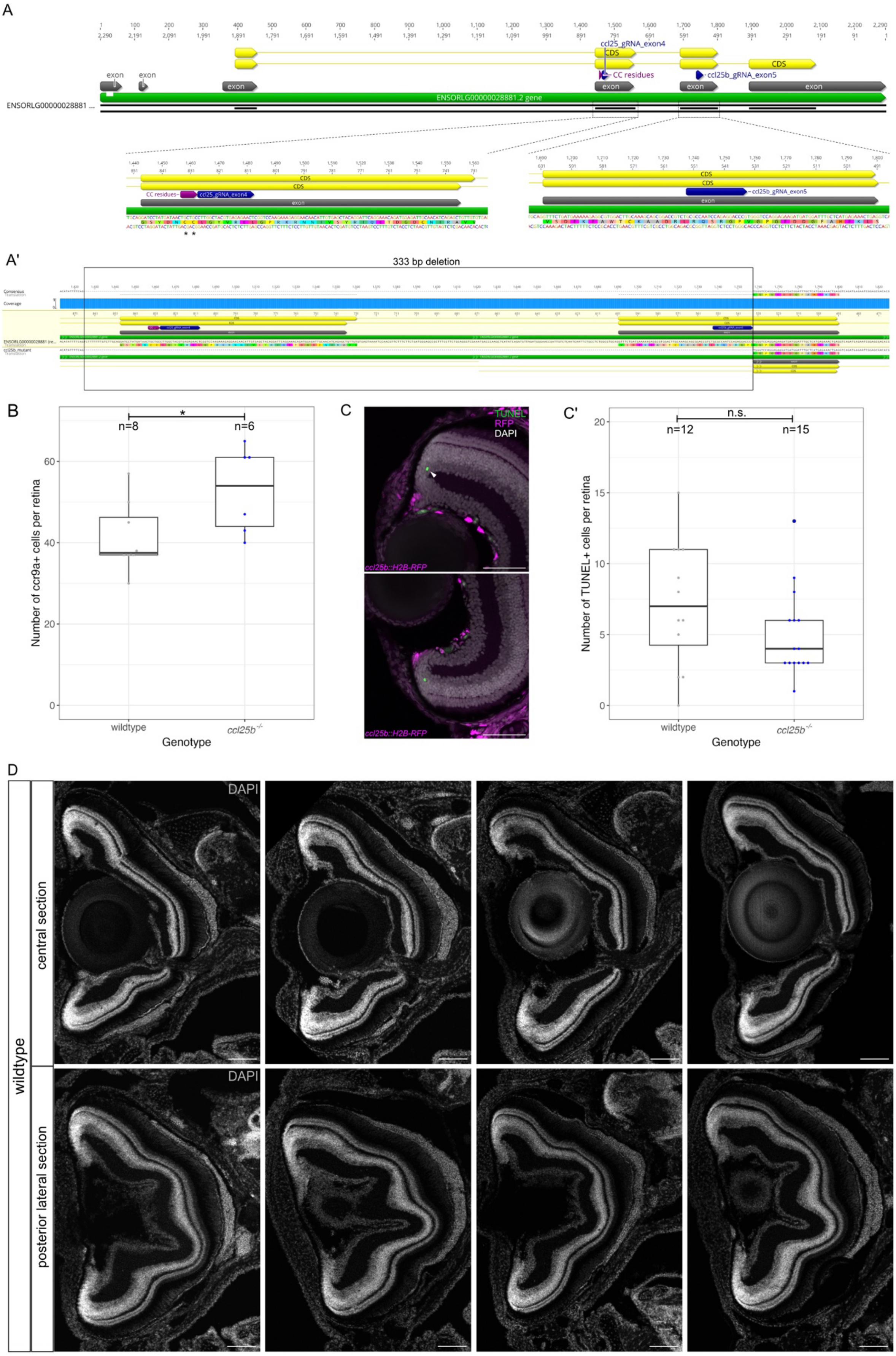
A
exon
exon
exon
ENSORLG00000028881.2 gene
ENSORLG00000028881 ...
ccl25_gRNA_exon4
CDS
CDS
CC residues
ccl25b_gRNA_exon5
A'
333 bp deletion
Consensus
Coverage
B
n=8
n=6
*
Number of ccr9a+ cells per retina
wildtype
ccl25b-/-
Genotype
C
TUNEL
RFP
DAPI
ccl25b::H2B-RFP
ccl25b::H2B-RFP
C'
n.s.
n=12
n=15
Number of TUNEL+ cells per retina
wildtype
ccl25b-/-
Genotype
D
wildtype
central section
posterior lateral section
DAPI
DAPI

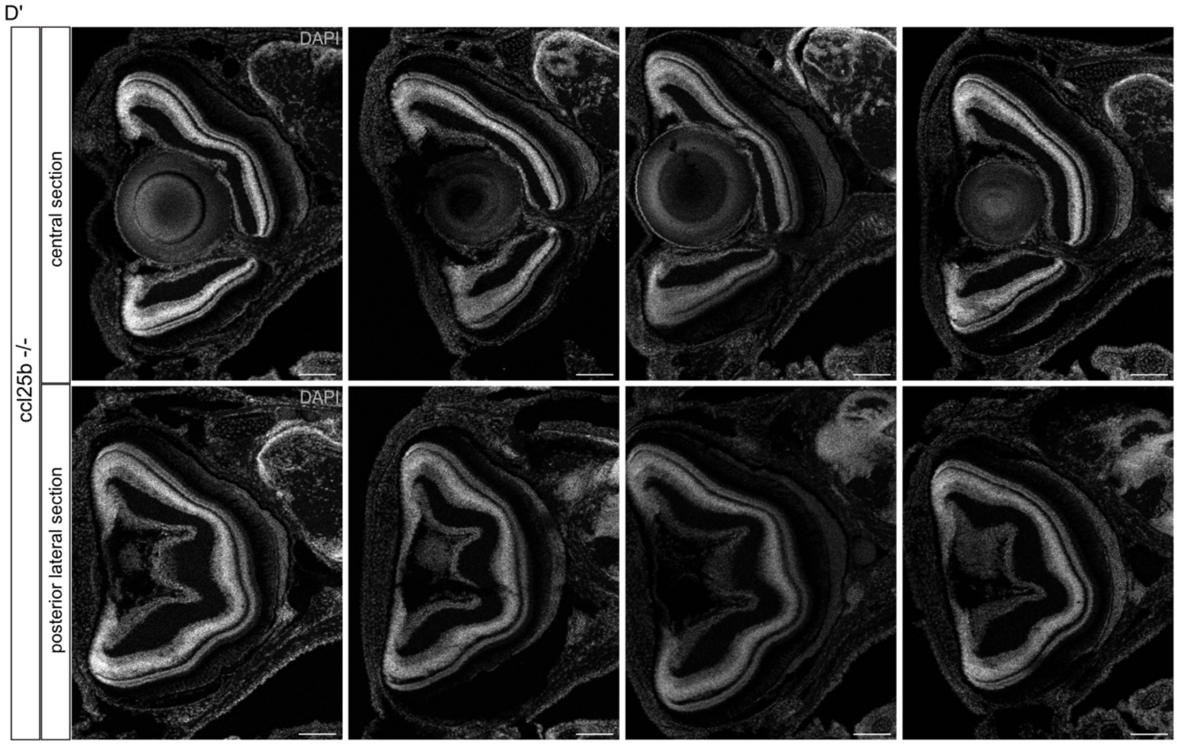


**Figure S3: Characteristics of *ccl25b* mutant**

(A) Genomic locus of the *ccl25b* gene consisting of 6 exons, including the CC motif in exon 4 and the gRNA target sites in exon 4 and 5 (zoomed regions). (A') Sequence alignment of the ccl25b locus comparing wildtype and *ccl25b*$^{-/-}$ mutant fish. The alignment reveals a 333 bp deletion in the mutant that eliminates exon 4 (containing the CC motif) and a portion of exon 5. (CC motif: magenta, coding sequence (CDS): yellow, exon: grey, gene locus: green, sgRNA: blue) (B) Quantification of *ccr9a*+ cells in retinae of *ccl25b*$^{-/-}$ (n=6) compared to wildtype (n=8) (*p = 0.04421; Mann-Whitney U test). (C) Apoptotic cells (green) detected in a transversal section of a *ccl25b::H2B-RFP* reporter hatchling retina by TUNEL staining. Scale bar: 50 µm. (C') Quantification of apoptotic cells in wildtype (n=12) and *ccl25b*$^{-/-}$ (n=15) retinae using TUNEL assay. (p = 0.2274; Mann-Whitney U test). (D-D') Overall morphology observed in transverse sections of central and posterior region wildtype and *ccl25b*$^{-/-}$ retinae stained with DAPI. Scale bar: 100 µm.

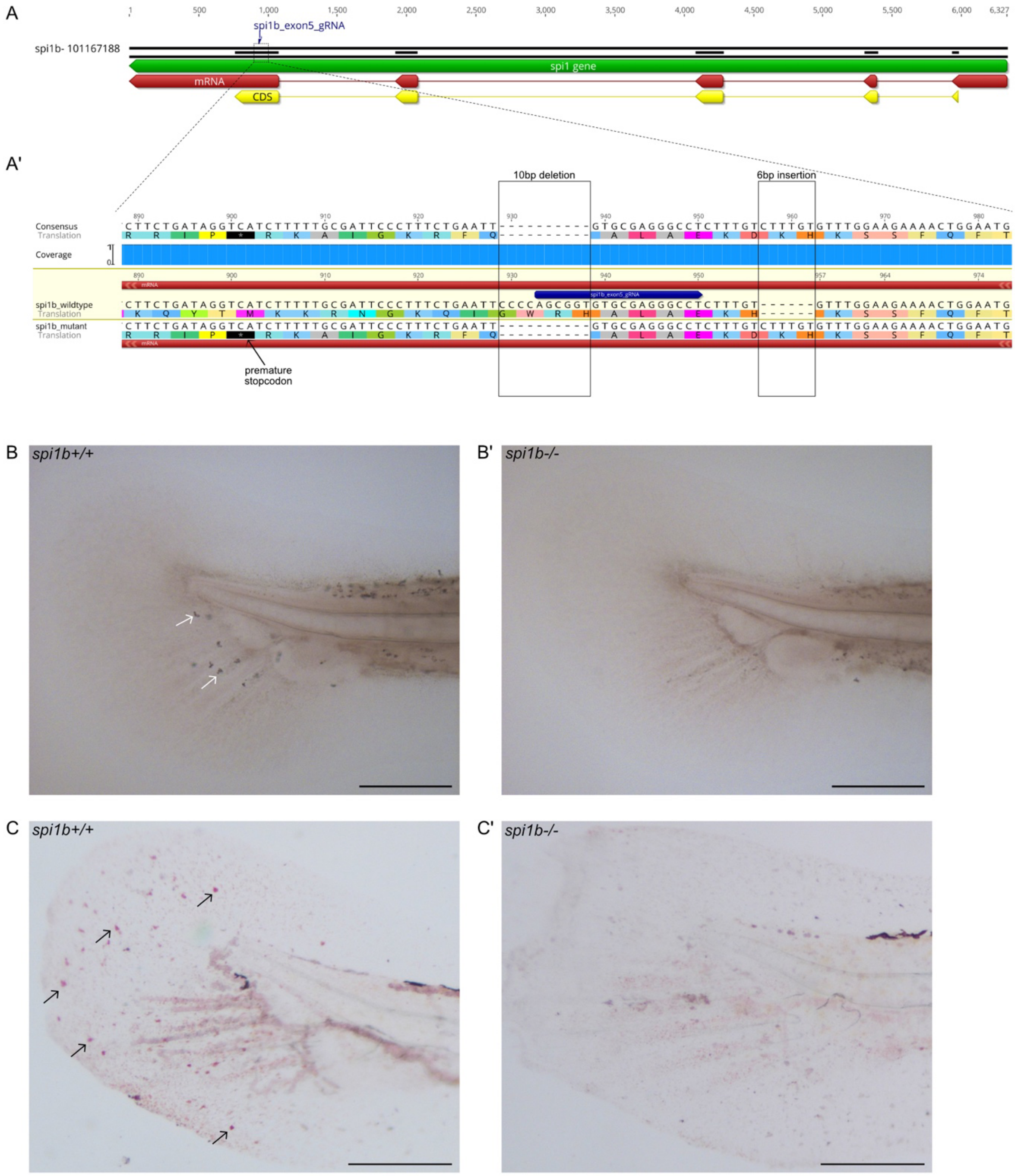
A
spi1b_exon5_gRNA
spi1b- 101167188
spi1 gene
mRNA
CDS
A'
10bp deletion
6bp insertion
Consensus
Translation
Coverage
spi1b_wildtype
spi1b_mutant
premature stopcodon
B spi1b+/+
B' spi1b-/-
C spi1b+/+
C' spi1b-/-

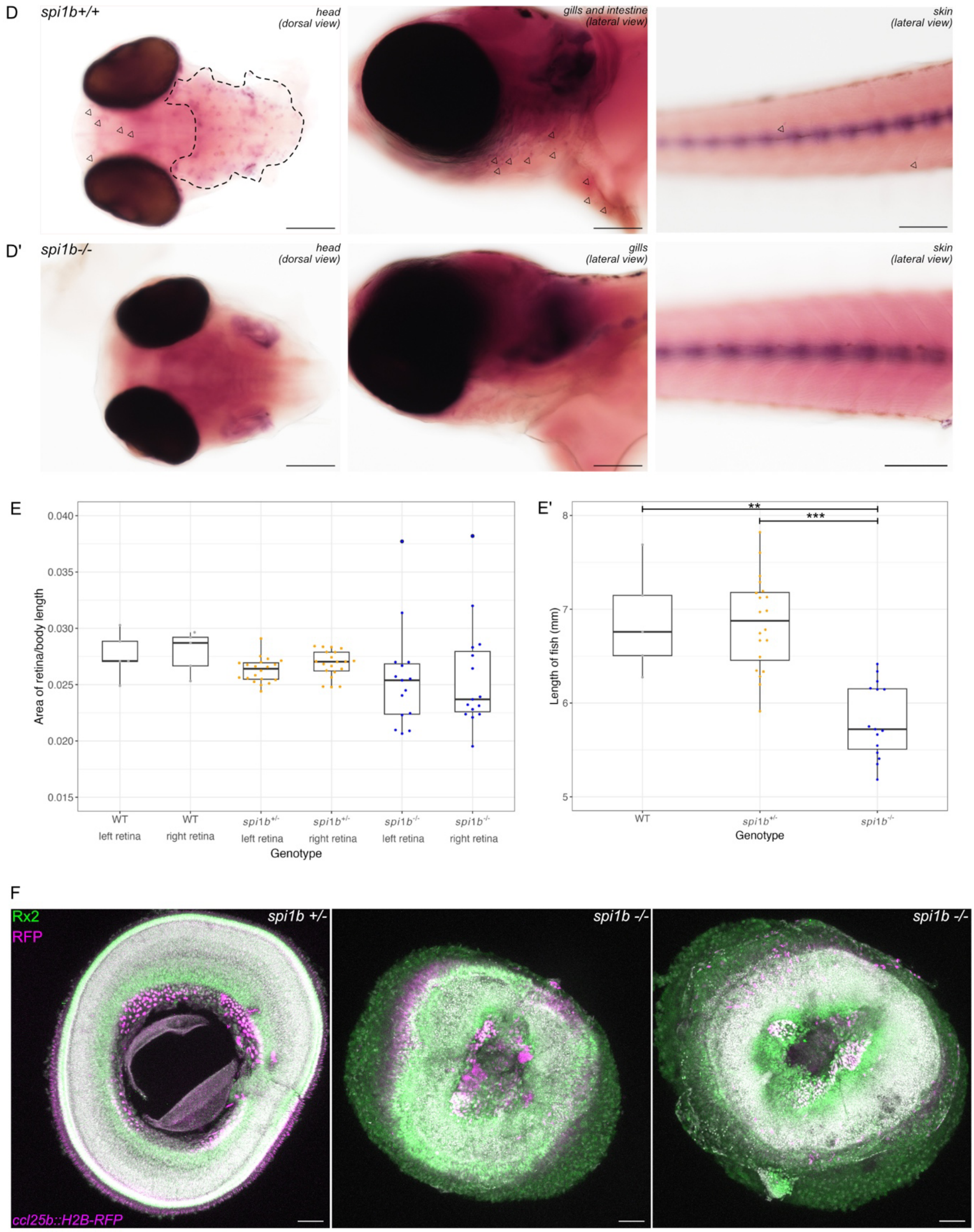


**Figure S4: The *spi1b* mutant lacks macrophages.**
(A) Overview of genomic locus of the *spi1b* gene showing location of 5 exons and the gRNA target site in exon 5 (green:gene; red:mRNA ; yellow:CDS). (A') Genetic alignment of the mutated region in the spi1b site with the WT sequence demonstrating a 10 bp deletion and 6bp insertion which leads to a frameshift and hence a premature stop codon (black box with asterisk) in the *spi1b* mutant. (sgRNA: blue, mRNA:red). (B) Sudan black staining to visualize

macrophages (white arrows) in WT and *spi1b*$^{-/-}$ hatchling tails. Macrophages in the tail regions (white arrows) are absent in *spi1b*$^{-/-}$ fish. Scale bar is 0.2 mm. (C-C') Neutral red staining in the tail fin of WT and *spi1b*$^{-/-}$ hatchlings to visualize presence of macrophages in *spi1b* mutant fish. Macrophages (black arrows) are present in the WT but not in *spi1b*$^{-/-}$ hatchlings. Scale bar is 0.2 mm. (D-D') In-situ hybridization of *cxcr3a* probe to visualize macrophages (black arrows and dotted line) in WT and *spi1b*$^{-/-}$ hatchling head, gill, intestine and skin of the posterior body. Macrophages are absent in *spi1b*$^{-/-}$ fish. (E) Ratio of left and right retina of WT (n=5), *spi1b*$^{+/-}$ (n=20) and *spi1b*$^{-/-}$ (n=15) hatchlings shows no significant difference in the eye sizes. (E') Length of WT (n=5), *spi1b*$^{+/-}$ (n=20) and *spi1b*$^{-/-}$ (n=15) hatchlings. ($p = 0.004281227$ for WT vs. *spi1b*$^{-/-}$, $p = 0.00001489453$ for *spi1b*$^{+/-}$ vs. *spi1b*$^{-/-}$, $p = 0.9590674$ for WT vs. *spi1b*$^{+/-}$ ; Dunn test with Holm-Bonferroni correction). (F) Whole mount immunostained retina comparing WT and *spi1b*$^{-/-}$ , show morphological defects in the *spi1b*$^{-/-}$ at 3 weeks post hatch.
